# The 1001G+ project: A curated collection of *Arabidopsis thaliana* long-read genome assemblies to advance plant research

**DOI:** 10.1101/2024.12.23.629943

**Authors:** The 1001 Genomes Plus Consortium, Carlos C. Alonso-Blanco, Haim Ashkenazy, Pierre Baduel, Zhigui Bao, Claude Becker, Erwann Caillieux, Vincent Colot, Duncan Crosbie, Louna De Oliveira, Joffrey Fitz, Katrin Fritschi, Elizaveta Grigoreva, Yalong Guo, Anette Habring, Ian Henderson, Xing-Hui Hou, Yiheng Hu, Anna Igolkina, Minghui Kang, Eric Kemen, Paul J. Kersey, Aleksandra Kornienko, Qichao Lian, Haijun Liu, Jianquan Liu, Miriam Lucke, Baptiste Mayjonade, Raphaël Mercier, Almudena Mollá Morales, Andrea Movilli, Kevin D. Murray, Matthew Naish, Magnus Nordborg, Fernando A. Rabanal, Fabrice Roux, Niklas Schandry, Korbinian Schneeberger, Rebecca Schwab, Gautam Shirsekar, Svitlana Sushko, Yueqi Tao, Luisa Teasdale, Sebastian Vorbrugg, Detlef Weigel, Wenfei Xian

**Author notes:** For correspondence: Magnus Nordborg and Detlef Weigel.

## Abstract

*Arabidopsis thaliana* was the first plant for which a high-quality genome sequence became available. The publication of the first reference genome sequence almost 25 years ago was already accompanied by genome-wide data on sequence polymorphisms in another accession, or naturally occurring strain. Since then, inventories of genome-wide diversity have been generated at increasingly precise levels. High-density genotype data for *A. thaliana*, including those from the 1001 Genomes Project, were key to demonstrating the enormous power of GWAS in inbred populations of wild plants, and the comparison of intraspecific polymorphism with interspecific divergence has illuminated many aspects of plant genome evolution. Over the past decade, an increasing number of nearly complete genome sequences have been published for many more accessions. Here, we highlight the diversity of a curated collection of previously published and so far unpublished genome sequences assembled using different types of long reads, including PacBio Continuous Long Reads (CLR), PacBio High Fidelity (HiFi) reads, and Oxford Nanopore Technologies (ONT) reads. This 1001 Genomes Plus (1001G+) resource is being made available at http://1001genomes.org. We invite colleagues with yet unpublished genome assemblies from *A. thaliana* accessions to contribute to this effort.

## Introduction

The human HapMap and 1000 Genomes Projects paved the way for the species-wide description of variation across the genome (The International Hapmap Consortium 2003; Birney and Soranzo 2015), but plants have not been far behind, with *Arabidopsis thaliana* being the second species for which genome-wide SNP data became available in 2007 (Kim et al. 2007). The same year saw the initiation of the 1001 Genomes Project for *A. thaliana*, an international, informal collaborative effort that demonstrated the practicality and power of resequencing large, publicly available collections of inbred lines that are perfect for genome-wide association studies (GWAS) (Atwell et al. 2010; 1001 Genomes Consortium 2016).

There was, however, a “dirty” secret underlying these all efforts, namely that the majority of genetic variation was largely ignored (Igolkina et al. 2024). While single nucleotide polymorphisms (SNPs) or small insertions and deletions (indels) are, with some caveats, accessible to short-read sequencing, the field has been mostly blind to major structural variants (SVs), such as large deletions, inversions, duplications, or polymorphic insertions of transposable elements (TEs). The importance of such major sequence differences has been documented in numerous functional studies (Weigel and Nordborg 2015; Alonge et al. 2020; Simon et al. 2022; Zhou et al. 2022; Jayakodi et al. 2024).

In response to the short-comings of short-read resequencing, several long-read technologies have been developed. In the last few years, dramatic advances in accuracy and cost of long-read sequencing have been made, allowing for population-scale analyses (De Coster et al. 2021). In *A. thaliana*, initial reports focused on individual genome sequences from non-reference strains, followed by larger and larger collections of long-read genome assemblies (Zapata et al. 2016; Jiao and Schneeberger 2020; Kang et al. 2023; Wlodzimierz et al. 2023b; Igolkina et al. 2024; Lian et al. 2024).

We describe ongoing efforts to generate a curated collection of *A. thaliana* genome sequences assembled from long reads, including both previously published and so far unpublished assemblies. We refer to this curated collection as the 1001 Genomes Plus (1001G+) resource. We are in the process of making these assemblies available at http://1001genomes.org, and we invite colleagues with collections of yet unpublished assemblies to contribute to this effort.

## Results

We collected published *A. thaliana* chromosome-level assemblies that were primarily generated with either long reads from Pacific Biosciences (PacBio) platforms (Continuous Long Reads [CLR], Consensus Circular Sequencing [CCS]/High Fidelity [HiFi] reads) or the Oxford Nanopore Technology (ONT) platform (Berlin et al. 2015; Chin et al. 2016; Zapata et al. 2016; Michael et al. 2018; Goel et al. 2019; Pucker et al. 2019; Jiao and Schneeberger 2020; Barragan et al. 2021; Naish et al. 2021; Wang et al. 2021, 2023; Hou et al. 2022; Rabanal et al. 2022; Wibowo et al. 2022; Christenhusz et al. 2023; Jaegle et al. 2023; Kang et al. 2023; Wlodzimierz et al. 2023b; Igolkina et al. 2024; Jiang et al. 2024; Kileeg et al. 2024; Lian et al. 2024; Tao et al. 2024; Teasdale et al. 2024). We also reached out to colleagues who we knew were still in the process of generating long-read assemblies. An overview of the number of genome assemblies from the different sources is provided in **Table 1**, and their diversity is indicated in **Figure 1**. In general, PacBio CLR-based assemblies reach excellent accuracy in the chromosome arms but are lacking most of the centromeres. PacBio HiFi-based assemblies have excellent accuracy and often span entire centromeres and 5S ribosomal RNA gene (rDNA) clusters within single contigs (Rabanal et al. 2022). ONT based assemblies have very good accuracy and include centromeric sequences, but these tend to be fragmented and do not span over chromosome arms. Finally, 45S rDNA arrays are very difficult to reconstruct completely; their assembly requires dedicated efforts that combine a variety of approaches (Fultz et al. 2023). The canonical chromosome number in *A. thaliana* is five, and many assemblies include telomeres for the eight chromosome arms that have non-repetitive sequences at their ends. The ends of two chromosome arms, 2p and 4p, consist of very large 45S rDNA arrays; telomeres can only be manually anchored to these two chromosome arms (Tao et al. 2024).

**Table 1.** Overview of *A*. *thaliana* genome assemblies in the 1001G+ collection.

| Techn. | Institution | Publ. | Unpubl. | References for published assemblies |
| --- | --- | --- | --- | --- |
| Legacy | Community | 1 |  | Lamesch et al. 2012 |
| PacBio CLR | Max Planck Institute (MPI) for Plant Breeding Research | 9 |  | Zapata et al. 2016; Goel et al. 2019; Jiao and Schneeberger 2020 |
|  | Bielefeld Univ. | 1 |  | Pucker et al. 2019 |
|  | MPI for Biology Tübingen | 2 | 5 | Barragan et al. 2021; Wibowo et al. 2022 |
|  | Gregor Mendel Institute of Molecular Plant Biology (GMI) | 6 | 135 | Jaegle et al. 2023 |
|  | University of Chinese Academy of Sciences | 2 |  | Wang et al. 2023; Jiang et al. 2024 |
|  | GMI/MPI for Biology Tübingen | 27 |  | Igolkina et al. 2024 |
|  | Institute of Botany, Chinese Academy of Sciences |  | 2 |  |
|  | Univ. of Tübingen |  | 16 |  |
| PacBio HiFi | Community |  | 2 | https://phoenixbioinformatics.atlassian.net/wiki/spaces/COM/pages/42215873/2024-01-15+PAG31+Summary |
|  | Darwin Tree of Life | 1 |  | Christenhusz et al. 2023 |
|  | MPI for Biology Tübingen | 72 | 87 | Rabanal et al. 2022; Wlodzimierz et al. 2023; Tao et al. 2024; Teasdale et al. 2024 |
|  | Lanzhou Univ. | 32 |  | Kang et al. 2023 |
|  | MPI for Plant Breeding Research | 48 |  | Lian et al. 2024 |
| ONT | Univ. of Cambridge | 1 | 1 | Naish et al. 2021 |
|  | Xi'an Jiaotong Univ. | 1 |  | Wang et al. 2021 |
|  | Univ. of Chinese Academy of Sciences | 1 |  | Hou et al. 2022 |
|  | MPI for Plant Breeding Research | 24 |  | Lian et al. 2024 |
|  | Univ. of Toronto | 8 |  | Kileeg et al. 2024 |
|  | Institut de Biologie de l'Ecole Normale Supérieure (IBENS) |  | 89 |  |
|  | Ludwig Maximilian Univ. Munich |  | 26 |  |
| **Total** |  | **238** | **361** |  |

**Figure 1.**
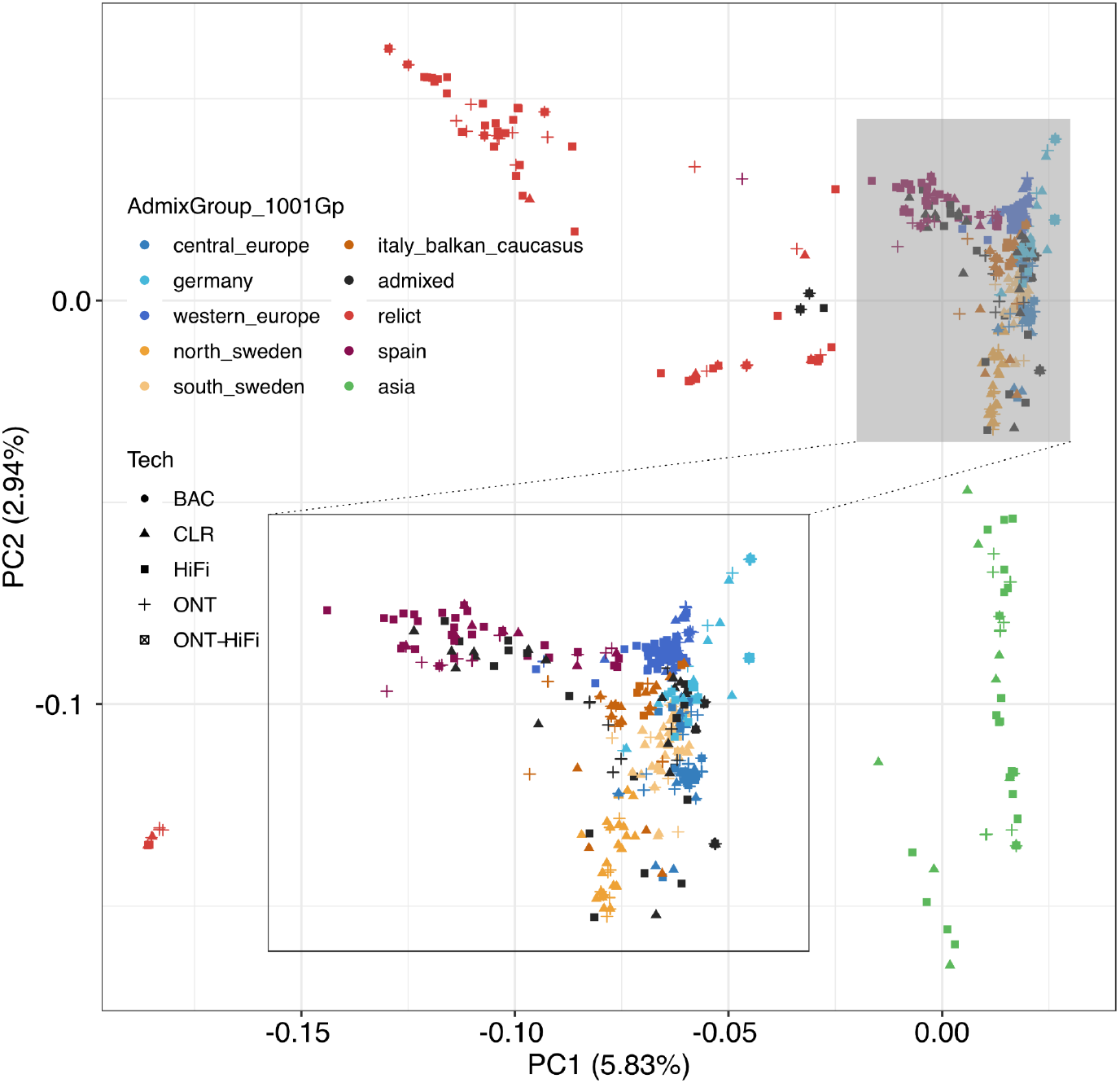
Diversity of *A. thaliana* assemblies. A Principal Component (PC) Analysis was performed based on bialleic SNPs from whole genome alignments of 581 assemblies to Col-CC (GCA_028009825.2, TAIR12). Colors indicate previously defined admixture groups (1001 Genomes Consortium 2016), shapes sequencing platforms. BAC, TAIR10 reference genome; ONT-HiFi, hybrid assembly from multiple Col-0 accessions. A few diverse assemblies, which are still being curated, are not yet included.

The assemblies are being made available in the Data Center of the 1001 Genomes project (https://1001genomes.org). None of the contigs in the original data sets are being reassembled, but we are performing quality checks for the scaffolding. Most of the available assemblies have been previously scaffolded based on reference genome information (Alonge et al. 2022), a process prone to errors. These errors often manifest as misplaced contigs, particularly those rich in repetitive sequences such as rDNAs and centromere satellite repeats, or organellar DNA contigs mistaken as true nuclear insertions (Rabanal et al. 2022). Another common error in publically available assemblies is the incorrect orientation of contigs in structurally variable regions. This is especially notable in the short arm of chromosome 4, where the reference accession Col-0 has a 1.7 Mb inversion spanning 1.17 Mb leading to the formation of a heterochromatic knob (Fransz et al. 2000), which is rare in the global *A. thaliana* population (Fransz et al. 2016). We therefore are visually inspecting all assemblies and using population information on structural variants to change contig joins that appear to be likely scaffolding errors due to reference bias. We are also identifying assemblies that are not from the accessions they are supposed to be based on previous short-read genotyping data, or that appear to have come from substantially identical material, sometimes with different accession identifiers. Finally, where original long reads are available, we are identifying regions of residual heterozygosity or potentially collapsed sequences in the assemblies. While the haploid assemblies are correct in the sense that the polymorphisms are present in the *A. thaliana* population, the heterozygous regions are reduced to a single haplophase that does not correspond to any haplotype that exists in the real world.

The total number of assemblies we have collected so far is 596 from 463 accessions. Minimizing also the number of accessions that are substantially related throughout their entire genomes (such as natural or lab-reared mutation accumulation lines (Exposito-Alonso et al. 2018; Monroe et al. 2022)), distinguished by fewer true SNPs than sequencing and assembly errors, the tally of unique assemblies currently stands at 438.

## Outlook

We are planning to release the complete set of curated assemblies in the second quarter of 2025. Analyses that the Weigel and Nordborg labs are currently conducting with these assemblies include the following:

- Annotation of nuclear sequences using liftoff (Shumate and Salzberg 2021), Helixer (Stiehler et al. 2021), and a custom pipeline that makes use of Iso-seq full-length cDNA data from 16 accessions (Teasdale et al. 2024).
- Assembly and annotation of plastid genomes (Xian et al. 2024).
- Annotation of telomere repeats (Tao et al. 2024).
- Annotation of ribosomal RNA genes and centromere satellite repeats (Wlodzimierz et al. 2023a).
- Annotation of TEs (Ou et al. 2019; Igolkina et al. 2024; Sierra and Durbin 2024).
- Analysis of synteny blocks (Wang et al. 2024).
- Annotation of short tandem repeats (Readman et al. 2021).
- Development of a JBrowse 2-based genome browser that allows for choosing any of the genomes as base for comparison with all other genomes (Diesh et al. 2023).

**We encourage colleagues who are generating additional genome assemblies of *A. thaliana* accessions or who are conducting either related or complementary analyses with available assemblies to join the 1001 Genomes Plus Project by contacting its coordinators, Magnus Nordborg and Detlef Weigel**. Similarly to the original 1001 Genomes Project, which was enormously successful despite never having received coordinated funding, we hope that the 1001 Genomes Plus Project can showcase the power of a federated, community-driven bottom-up approach to the generation of powerful resources for evolutionary genomics.

## Methods

### Variant calling

We aligned 581 *A*.*thaliana* and two *A. lyrata* genome assemblies (Kolesnikova et al. 2023) to the Col-CC reference genome (GCA_028009825.2, TAIR12) using wfmash (v0.13) with parameters “-s 10k -p 90 -n 1”. Variants were called across multiple samples with minipileup (https://github.com/lh3/minipileup, v1.0) using “-s0 -a0 -q0 -l 20000 -vcC -f Col-CC.fa”. Variants in centromeric and rDNA regions were excluded using bcftools (Danecek et al. 2021). Genomes were also aligned to TAIR10, and genotype IDs were verified by scores >0.98 with SNPmatch (Pisupati et al. 2017) and the 1001G callset as the database.

### Population genetics analysis

We created two biallelic SNP subsets: one including *A. lyrata* as an outgroup for a neighbor-joining (NJ) tree, and the other for kinship and principal component analysis (PCA). For the NJ tree, we included sites where both *A. lyrata* accessions shared the same allele and filtered for a minor allele frequency (MAF) >0.01 and a missing rate <20%. Distances were calculated with VCF2Dis (https://github.com/BGI-shenzhen/VCF2Dis, v1.50), and the tree was constructed using fneighbor in PHYLIP (Felsenstein 1989). For PCA and kinship analyses, we excluded *A. lyrata* and applied the same filtering thresholds. PCA was conducted using GCTA (Yang et al. 2011) (v1.94), retaining the first 20 principal components, and related individuals were identified with KING (Manichaikul et al. 2010) (--related). Admixture groups were assigned based on clustering in the neighbor-joining tree, with missing labels resolved through majority voting among labeled samples from SNPmatch-based IDs within each clade.

## Acknowledgments

We thank all members of the global *A. thaliana* community who have directly or indirectly contributed to the collection of germplasm and the generation of genome sequences. Supported by grants PID2022-136893NB-I00 from the MCIN/AEI/10.13039/501100011033 and FEDER (EU) (Carlos Alonso-Blanco), TRR356 (DFG 491090170) (Niklas Schandry), ANR STEVE (ANR-21-CE45-0018) (Vincent Colot), the Austrian Academy of Sciences (Magnus Nordborg), the Novozymes Prize of the Novo Nordisk Foundation and the Max Planck Society (Detlef Weigel). The initial phase of the 1001 Genomes Plus Project in the Kersey, Nordborg and Weigel labs was supported by DFG, FWF and BBSRC through ERA-CAPS grant 1001G+.

